# Is the radial growth of irrigated urban trees more strongly correlated to light and temperature than water?

**DOI:** 10.1101/2020.10.29.361121

**Authors:** Kevin L. Griffin, Thomas G. Harris, Sarah Bruner, Patrick McKenzie, Jeremy Hise

**Author notes:** **Corresponding Author**: Kevin L. Griffin, Lamont-Doherty Earth Observatory of Columbia University, 61 Route 9W, Palisades NY 10964. (845)422-1364.

## Abstract

**Background:** Real-time monitoring of tree growth can provide novel information about trees in urban/suburban areas and the myriad ecosystem services they provide. By monitoring irrigated specimen trees we test the hypothesis that in trees with sufficient water, growth is governed by environmental factors regulating energy gain rather than by factors related to water use.

**Methods:** Internet enabled, high-resolution dendrometers were installed on three trees in Southampton, NY. The instruments, along with a weather station, streamed data to a project web page that was updated once an hour. (https://ecosensornetwork.com). Growing periods were determined using a Hidden Markov Model based on Zweifel et al.’s (2016) zero-growth model. Linear models and conditional inference trees correlated environmental variables to growth magnitude and rate of growth.

**Results:** Growth was governed by the interacting environmental variables of air temperature, soil moisture, VPD and took place primarily at night. Radial growth of spruce began April 14 after the accumulation of 69.7 °C growing degrees days and ended September 7th. Cedar growth began later (April 26^th^), after the accumulation of 160.6 °C and ended later (November 3^rd^). During the observation period, these three modest suburban trees sequestered 108.3 kg of CO_2_.

**Conclusions:** Though irrigated, residential tree growth in our experiment was affected by environmental factors relating to both water use and energy gain through photosynthesis. Linking tree growth to fluctuations in environmental conditions facilitates the development of a predictive understanding useful for ecosystem management and growth forecasting across future altering climates.

## Introduction

The biogeophysical environment of urban areas is substantially different from that of rural ones. Humans are altering the hydrologic cycle of exurban watersheds, both by importing vast amounts of fresh water (Booth & Bledsoe, 2009) and through the widespread use of impervious materials that modify the local water balance by decreasing groundwater recharge and increasing surface water runoff (Haase, 2009; Walsh et al., 2012). Urban centers modify the local climate by releasing sensible heat from artificial surfaces warmed during the day (Weng, 2003; Hamin & Gurran, 2009; McGeehin & Mirabelli, 2001). Airborne particulates in urban areas can create rain-inducing condensation nuclei that result in increased precipitation in, and downwind of, cities (Shepherd et al., 2002). Urban air itself contains increased concentrations of pollutants such as CO_2_, nitrogen oxides, sulfur oxides, ozone and other volatile organic compounds (Chameides et al., 1992; Idso et al., 1998; Lovett et al., 2000; Nowak et al., 2006; Calfapietra et al., 2013). Urban soils often contain elevated levels of heavy metals (Bernhardt et al., 2008; Kaushal & Belt, 2012; Sonti et al., 2019). Furthermore, changes in rates of litter decomposition (Carreiro et al., 1999), mineralization and nitrification (Zhu & Carreiro, 2004), and shifts in soil faunal communities (Steinberg et al., 1997) affect local biogeochemical cycles of essential nutrients. Finally, increased abundance of exotic species in urban areas is contributing to significant changes in species composition in urban ecosystems (Rudnicky & McDonnell, 1989). Understanding how these biogeophysical conditions and changes impact urban ecosystem services and the communities that depend on them is of critical importance.

Trees and tree growth provide a multitude of important benefits for urban ecosystems and their residents that can help mitigate many of the negative changes caused by urbanization (Figure 1). Among the many commonly recognized ecosystem services (e.g. Nitoslawski et al., 2019) provided by urban forests are: carbon sequestration and storage (Nowak & Crane, 2002; Pataki et al., 2006; Edmondson et al., 2012); stormwater and flood water management (Denman et al., 2016); improved nutrient cycling (Livesley et al., 2016); local cooling (Norton et al., 2015; Solecki et al., 2005); reductions of air pollution (Dochinger, 1980; Nowak et al., 2006; Abhijith et al., 2017); and increases in biodiversity (Ikin et al., 2013). At the same time, tree function, e.g. growth, reproduction, and survival, is heavily influenced by these same environmental factors. In this way, trees play a key role in urban ecosystems both affecting and being affected by the biogeophysical environment and thus understanding the physiological underpinnings of tree growth in urban forests is essential.

**Figure 1.**
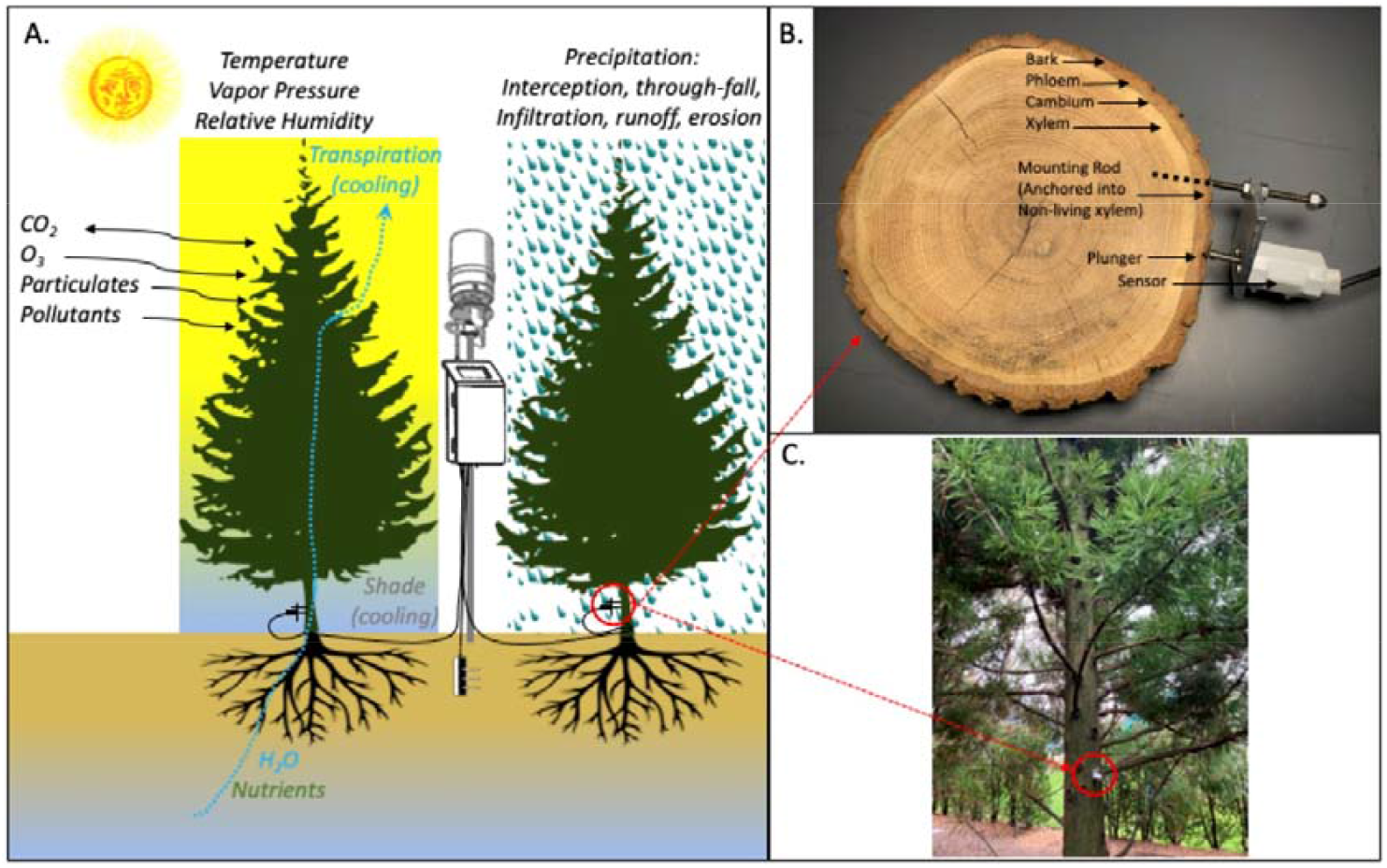
Conceptual overview of the environmental drivers of urban tree growth and resulting ecosystem services provided by suburban trees considered when monitoring tree growth with point-dendrometers. A) Experimental protocol relies on point dendrometers attached to tree stems at 1.37 m from the ground. Also shown are the all-in-one weather station, soil temperature and moisture sensor and cell phone enabled data logger. Urban trees provide a variety of ecosystem services including: local cooling, carbon sequestration, removal of air pollutants, reduced erosion and runoff, and improved nutrient cycling. B) A dendrometer is mounted to the heartwood and holds the sensors in contact with the bark. The measurement detects changes in the elastic tissues below including the bark, phloem, cambium, and immature xylem. C) a photograph of one of the study trees highlighting the installed dendrometer.

Traditionally tree growth is measured infrequently at low resolution. For example, foresters use methods of the USDA -Forest Service’s Forest Inventory Analysis program (https://www.fia.fs.fed.us) such as recording diameter at breast height (DBH = 1.37m from the ground) to calculate basal area. Assessed yearly, these can provide growth rates; tree cores yield similar information and can be taken at a single time point, but require additional analysis (Sonti et al., 2019). Street tree surveys of size and health are part of many urban parks programs (Nowak et al., 2018) and aerial photographs (Spurr & Brown Jr, 1946) can provide useful insights as to quality and distribution of urban greenery. More recently, technological solutions such as aerial LiDAR (Alonzo et al., 2014), drones (Elliott, 2016), and satellites (Palomino et al., 2017; reviewed in Nitoslawski et al., 2019) show great potential to add in the understanding of tree growth in urban environments. All of these approaches, however, require repeat observations which can be expensive and/or infrequent. As a result of the limited spatial and temporal scales of the tree growth products typically obtained, our understanding of the links between the biogeophysical environment and the physiological underpinnings of tree growth in urban forests is similarly limited.

Dendrometers provide an alternative method of monitoring radial growth of trees and could provide much needed high-resolution data in urban ecosystems. When mounted to the non-living sapwood/heartwood of the tree, these highly responsive tree growth sensors can record when new cells are produced by the adjacent vascular cambium and/or the hydration of the xylem, cambium, phloem, or bark (Zweifel et al., 2014). Not only can these instruments provide detailed records of radial stem growth and water use (Steppe et al., 2006; Zweifel et al., 2006), when used in urban settings the instruments are easily connected to the internet and can stream data live in real or near-real time. The result is a time series of stem radius that can be used for research, education and outreach and provides a much-needed link between growth and the environment.

In this study, we installed three dendrometers in a suburban backyard along with a weather station to study the short time-scale interactions between tree growth and meteorological conditions in urban environments. We use a time series spanning 1.5 years to model tree growth and the relationship to local weather and climate. We define statistical relationships between growth, and light, temperature, soil moisture and growing degree days to test the hypothesis that, in this urban setting where trees are irrigated, diameter growth is more sensitive to environmental variables related to energy gain (light and temperature) than to factors related to water use (precipitation, soil moisture and the atmospheric Vapor Pressure Deficit (VPD). We also discuss the usefulness of this monitoring approach and the research, education and outreach potential of an expanded dendrometer network in urban forests.

## Methods

### Location and Plant Material

Three adjacent ornamental trees growing in a backyard tree in Southampton, NY (USA), were used for this study, two white spruce (*Picea glauca* (Moench) Voss) and one Japanese cedar (*Cryptomeria japonica* (L. f.) D. Don). The experiment was established in June of 2018. The native soils of this location are of the Bridgehampton silt loam association (NRCS, 2009) and are both well-drained and inherently low in mineral nutrition. Significant soil improvement, modification and the addition of organic materials and fertilizers has taken place. The study area is irrigated and thus the study trees rarely experience prolonged water limitation. The climate of Southampton is maritime and strongly seasonal with historical July (the warmest month) minimum and maximum temperatures averaging 18 and 27 °C respectively and historical January (the coldest month) minimum and maximum temperatures averaging -4.4 and 3.9 °C respectively (https://prism.oregonstate.edu/explorer/). Rainfall is evenly distributed throughout the year averaging 106.43 mm month^-1^ and totaling 1.28 m yr^-1^. Growing degree days were also calculated as the accumulated average temperatures above 5 °C following the coldest day of the year.

### Diameter Growth and Environmental Monitoring

A time series of tree diameter was recorded by a point dendrometer (Hise Scientific Instrumentation, Somers, New York USA). The dendrometer is attached to the stem of the tree using a mounting bracket that maintains the potentiometer in a fixed position relative to the non-living xylem with a stainless steel 6.35 mm diameter rod, inserted into the stem. All dendrometers were placed on the north side of the tree (to minimize exposure to direct solar radiation) at a height of 1.37 m from the ground surface. Dendrometers were installed on top of the thin bark (after carefully removing any loose bark with a small rasp, while avoiding damage to the living tissues), and thus all processes causing the changes in stem diameter below the location of the sensor were recorded. These may include: bark swelling and shrinking, cambial activity, changes in phloem, xylogenesis and stem hydraulics. Every five minutes the average position of the sensor was recorded (EM50G, METER Group Inc, Pullman, WA USA). The installation method and hardware have been extensively tested against other commercially available dendrometers and were found to be robust with no significant relationships between air temperature, relative humidity (RH), VPD, wind speed, wind direction or wind gusts and sensor output (Kendall’s Tcoefficient 0.039, 0.003, -0.207, 0.095, -0.177, 0.047, respectively; all *p*<0.001). While the single point of attachment could result in disturbance by animals these would be easily detectable as distinct rapid changes in diameter unrelated to observed trends in the biology of the stem or the current ambient environmental conditions and therefore could be easily identified and removed from the data. No such disturbances were detected. Furthermore, by making only a single penetration of the stem, offset from the column of vascular tissue influencing the water transport beneath the sensor, we feel this mounting system has some potential advantages over other multipoint attachment methods. Each potentiometer was independently calibrated at the time of installation and output is considered linear over the very small annual change (<3 mm). Each sensor’s initial voltage was related to the measured stem diameter at DBH, which allowed us to continuously calculate the stem radius.

Local environmental conditions at the experimental site were monitored throughout the experiment using a digital weather station (ATMOS 41, METER Group Inc, Pullman, WA USA). This station measures: Solar radiation, precipitation, air temperature, barometric pressure, vapor pressure, relative humidity, wind speed, wind direction, maximum wind gust, lightning strikes, lightning distance and tilt. In addition, soil moisture, soil temperature and conductivity were measured in a single location, at a depth of 10 cm, approximately 2.5m from Spruce 2 and Cedar (TEROS 12, METER Group Inc, Pullman, WA USA). All environmental, soil, and growth data were recorded as five minute averages with a single data logger (see above) and each hour these data were automatically uploaded to the Zentra Cloud (METER Group Inc, Pullman, WA USA). From here the data were transferred to the Eco Sensor Network (ESN, Hise Scientific Instrumentation, Somers, New York USA) where the data are publicly available in near real time (https://ecosensornetwork.com).

To convert changes in stem radius to biomass, biomass increments and rates of aboveground carbon sequestration, standard geometric relationships were assumed so that allometric biomass equations could be applied to the calculated DBH (stem diameter in cm measured 1.37m from the ground). White spruce aboveground biomass (kg) was calculated as 0.0777*DBH^2.4720^ (Harding & Grigal, 1985). No allometric equation specific to *Cryptomeria japonica* was found so biomass was calculated as exp(−2.0336 + 2.2592*ln(DBH)), a general equation for cedar and larch (Jenkins et al., 2003). Root mass for both species was conservatively estimated at 30% of the aboveground mass, based on the results of white spruce total biomass harvest performed by Ker and van Raalte (1981). Carbon sequestration was calculated as the increment in dry mass accumulation, assuming the carbon content of woody biomass is 50% (Thomas & Martin, 2012). By extension CO_2_ can then be calculated based on the atomic ratio of one carbon atom to two oxygen atoms in each molecule of CO_2_ with atomic weights of 12 and 16 grams per mole (a total of 44 grams per mole of CO_2_), thus the weight of carbon was multiplied by 3.67.

### Statistical analysis

R version 3.4.4 was used for all statistical analyses (R Development Core Team, 2006). Due to occasional low battery levels, not every 5-minute interval was recorded during a 24-hour period. Days missing more than 25% of their data were removed.

### Growth Models

Growing seasons for each tree were determined by using a piecewise regression which iteratively searched for a breakpoint by minimizing the mean squared error of the segments (Crawley, 2012). This was done separately for each spruce at the end of each season, and the cedar at the beginning and end of each season. The beginning of the growing season for the spruces were determined by the day they reached the previous year’s maximum stem radius. These individual growth seasons were used for subsequent statistical analysis.

A zero-growth model (ZG, Zweifel et al., 2016), where “growth” began when the current dendrometer exceeded the last maximum and ended at the first decreasing reading, determined periods of irreversible radial growth within the growing season. Since the ZG model is sensitive to any change in the dendrometer reading, it has the potential to overestimate the number of starts and stops in growth, resulting in a very high number of five minute growth periods (the recording interval of the data logger). We used a Hidden Markov Model (HMM) to summarize the ZG model intervals and make them more suitable for statistical analysis. The HMM assigned each time point to one of two latent variable states: “growth” or “deficit” (Durbin et al., 1998; Eddy, 2004). The model was trained using emitted states of the Zweifel et al. (2016) ZG model. We used the Baum-Welch algorithm to infer model parameters from the full data for each tree, and we used the Viterbi algorithm to infer the most likely state of the latent variable at each point (Durbin et al., 1998). All HMM code was implemented using the package ‘HMM’ in R (R Development Core Team, 2006; Himmelmann, 2010).

For downstream statistics, we isolated the HMM-identified “growth” periods that overlapped with more than two data points. From these growth periods we derived our response variables and our predictors. The response variables are 1) the net change in the zero-growth model through the growth interval (“total growth;” y_end_-y_start_) and 2) the value from (1) divided by the duration of the growth interval (“slope”). The value of (2) corresponds to the slope of a line passing through the zero growth model values at the beginning and end of the growth interval. The predictors are environmental variables. We calculated their values using the mean through the growth interval.

### Linear models

As most of growth occurs at night, we conducted a multiple regression analysis on the HMM-identified growth periods when solar radiation was equal to zero. The models were applied separately to each tree and to the total growth and slope response variables for each interval. During each interval, we used the mean of each environmental factor and their interactions as the explanatory variables.

### Conditional inference trees (CITs)

While the linear models test for the effect of a variable across all of the data, interactions between variables could result in cases where a particular explanatory variable only has a significant influence over the response in a particular part of another variable’s range. With this in mind, we fit conditional inference trees (CITs) to further examine the association between the explanatory and response variables. In comparison to the linear models, which tested the influence of each variable across the full data, the CITs repeatedly partitioned the data using the predictors. For example, even though “variable X” might not show an influence on the whole data --so a linear model would show no significance --“X” might instead have a significant influence for the subset of the data where a different “variable Y” has a high value.

The CITs were fit on the HMM-identified growth periods for each tree, with response variables being the “total growth” and “slope” metrics defined in the previous section. The predictors were the mean values of the environmental variables through each interval. We performed CIT analysis using the *partykit* package in R (Hothorn & Zeileis, 2015).

## Results

### Micrometeorological conditions

Air temperature during our experiment varied from a low of 4.8°C (5/1/19) to a high of 32.0 °C (7/14/19), with a mean temperature of 19.37± 5.18 °C (Figure 2). Daytime temperatures were on average 3 °C degrees warmer than nighttime temperatures, 20.6 ± 5.15 and 18.0±4.84 respectively, during the experimental period. Soil temperatures varied between 24.4 (8/9/18) and 8.7 with a mean of 18.3 ± 4.26 °C (Figure 2). Total precipitation was 1012 mm, coming in 28 discrete events averaging 18 mm (Figure 2). The combination of rain and irrigation resulted in a mean volumetric soil water content of 0.38±.026 m^3^ m^-3^. The wind was primarily onshore, coming from the south/southwest (201 °) (data not shown). The total solar energy received during the experimental period was 8.9 MW m^-2^, with a daily average of 816.6 W m^-2^ d^-1^.

**Figure 2.**
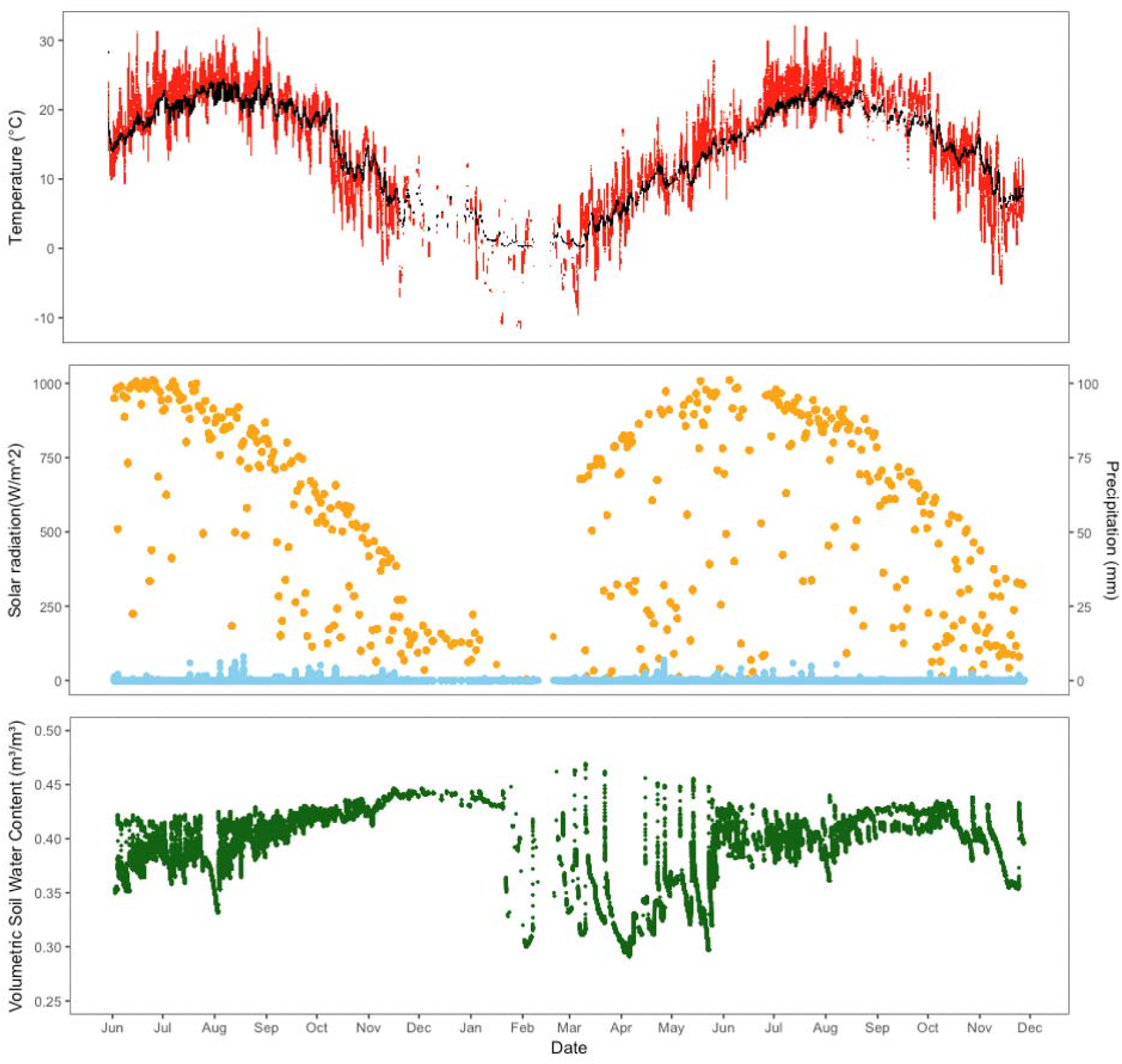
Weather conditions over the two-year period of the study (2018 to 2019).. Maximum daily soil temperature (black) and air temperature (red) (□) are in the top panel, solar radiation (orange) (W m^-2^) and precipitation (blue)(mm) are in the second panel, and Volumetric Soil Water Content in the third (mm mm^-3^).

### Stem diameter

The study trees were of average size in this planted landscape, similar to most of other trees within a 100m. Over the duration of the experiment, these trees grew by an average of 11% reaching final diameters of 13.4, 15.9 & 17.34 (Table 1, Figure 3). In 2019, the period of rapid diameter growth began on 4/14 ± 10 days, and ended on 9/07 ± 10 days (spruce) and began on 4/26, and ended on 11/3 ± 5 days (cedar, dotted vertical lines on Figure 3). Note a small portion of the 2018 early season growth may have occurred prior to the start of this experiment and thus be absent record and the two white spruce trees grew enough to saturate the dendrometer signal by end of the 2019 growing season (Figure 3a & b). Across the growing seasons, the ZG model (expected to be very sensitive to short-term fluctuations) identified 6096 growth periods (mean duration 13.6 minutes, standard deviation 15.4 minutes), and the HMM results then consolidated these to 843 growth periods (mean duration 152.5 minutes, standard deviation 205.4 minutes). The basal area increments added to these trees were 37.2, 29.0 and 7.8 cm^2^ (for Spruce 1, Spruce 2 and Cedar respectively) during the experiment.

**Table 1.** Characteristics of the four study trees, including Diameter at Breast Height (DBH), Basal Area Increment (BAI), and biomass. Above ground biomass (AGB) estimated using allometric equations. Below ground Biomass estimated as 30% of AGB, see text for more details.

|  | DBH (cm) | BAI (m2) | Above Ground Biomass |  |  | Below Ground Biomass |  |  |
| --- | --- | --- | --- | --- | --- | --- | --- | --- |
|  |  |  | Initial (kg) | Final (kg) | Gain (kg) | Initial (kg) | Final (kg) | Gain (kg) |
| Spruce 1 | 11.5 | 37.2 | 33.3 | 48.4 | 15.1 | 8.2 | 11.4 | 3.1 |
| Spruce 2 | 14.0 | 29.0 | 54.5 | 70.5 | 16.0 | 12.6 | 15.7 | 3.1 |
| Cedar 1 | 16.7 | 17.3 | 77.0 | 91.3 | 14.4 | 18.0 | 21.1 | 3.1 |
| Total |  | 83.50 | 164.81 | 210.19 | 45.39 | 38.83 | 48.07 | 9.25 |
| Mean |  | 27.83 | 54.94 | 70.06 | 15.13 | 12.94 | 16.02 | 3.08 |
| SEM |  | 5.774 | 12.596 | 12.395 | 0.471 | 2.827 | 2.805 | 0.025 |

**Figure 3.**
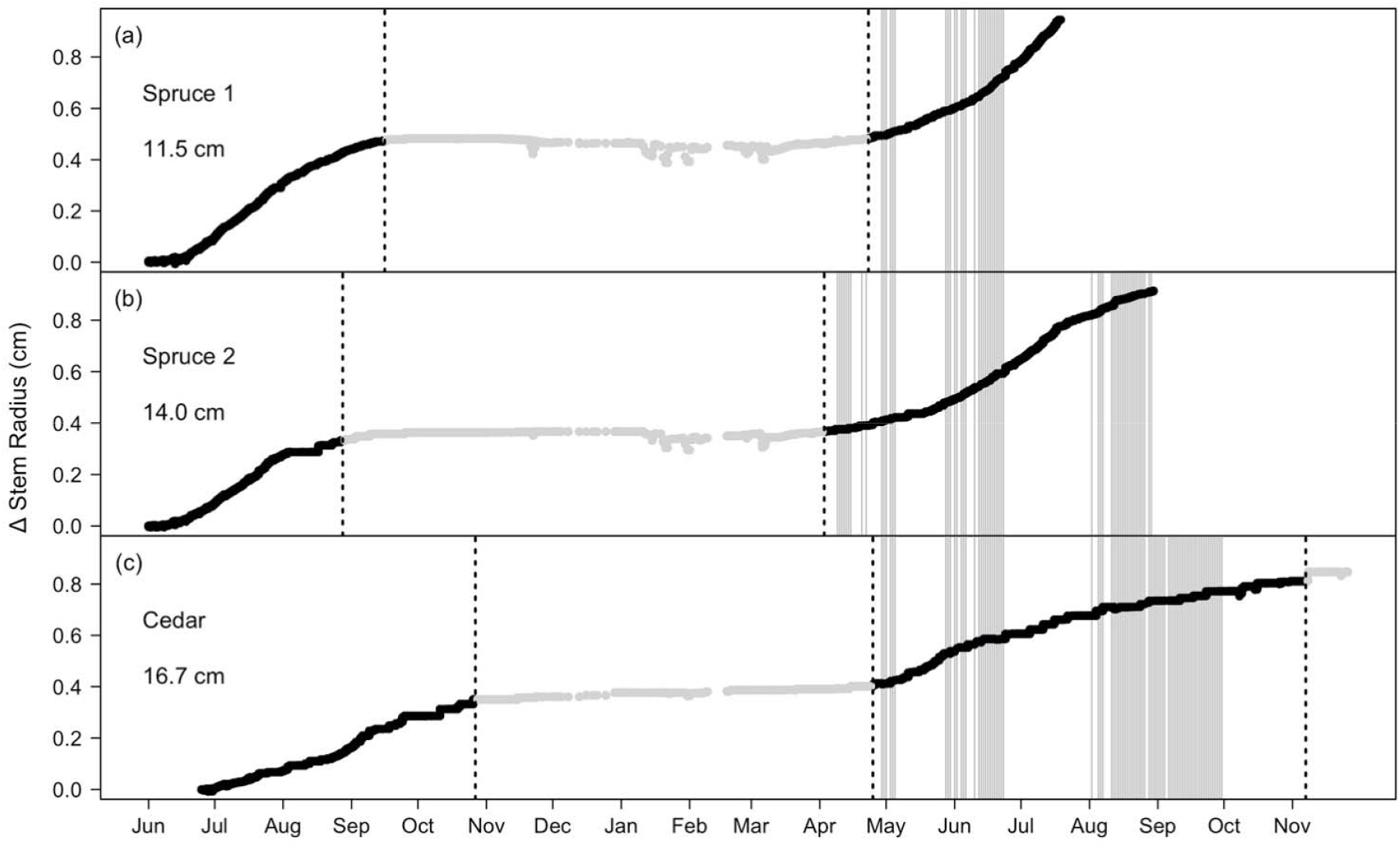
Tree Growth, measured as linear distance (cm) from June 03 2018 until November 11 2019. Replicate trees of spruce (*Picea glauca*, a and b) and a cedar (*Cryptomeria japonica*, c) are shown along with each tree’s initial diameter (cm) at the height of the sensors. Dotted vertical lines show the calculated end of the 2018 growing season, start of the 2019 growing season, and end of the 2019 growing season (cedar only). The end of the 2019 spruce signal is related to reaching the maximum sensor displacement. Lines highlighted in grey indicated loss of 25% or more of daily data as a result of battery life.

### Growth with and without hydrologic constraints

Figure 4 displays two seven-day periods: June 12th-19th, a period of limited growth, but large hydrologic fluctuations (Figure 4a, c, e & g) and July 14th-21th, a period of consistent stem growth (Figure 4b, d, f & h). The ZG model reveals only a single period of water deficit during the period of consistent growth (Figure 4f & h), which coincides with higher VPD and declining soil moisture (Figure 4d). Over this interval, some growth was experienced on each day, although this was minimal on July 19 following the only instance of water deficit. During the period of limited growth, the ZG model shows three intervals of substantial growth (June 12th-14th), followed by an extended time of water deficit, reaching a maximum on June 14th and continuing until June 18th. Growth periods as differentially identified by the ZG and HMM models are shown across panels e-h as vertical grey stripes.

**Figure 4.**
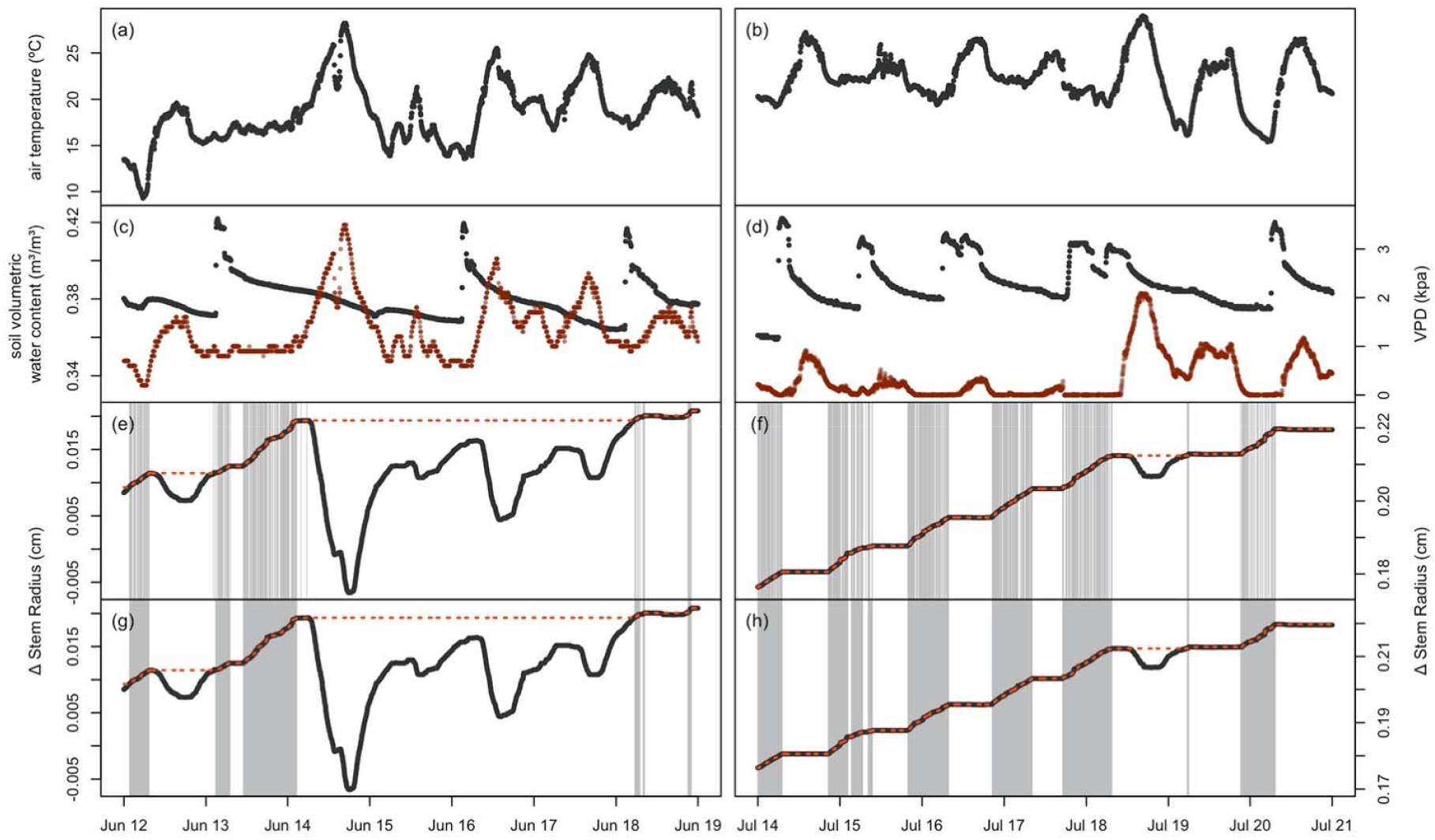
Environmental variables and radial growth over two, one-week time periods, one with a stem experiencing dehydration (6/12/18 to 6/19/18) and one hydrated stem (7/14/18 to 7/21/18). a) Air temperature for dehydrated period and b) hydrated period. c) Soil moisture (black points) and vapor pressure deficit (VPD, brown points) for dehydrated period and d) hydrated period. e) dendrometer output (black line), zero-growth model (ZG, red dashed line), and ZG model identified growth periods (grey vertical stripes) for dehydrated period and f) hydrated period. g) dendrometer output (black line), ZG model (red dashed line), and HMM identified growth periods (grey vertical bars) for dehydrated period and f) hydrated period. Note that the two time periods use different scales for the dendrometer output as significantly more growth occurs during the hydrated period.

### Linear Models

We used multiple regression models to identify which combination of environmental variables had the most predictive power estimating growth (Table 2). The rate of growth for all trees was best explained by a model including air temperature, soil moisture, VPD, and all of the interaction terms. No single environmental variable was significant on its own. This model explained slope for all trees, but only growth (the magnitude of expansion during the growth periods) for the spruce trees. While the model was significant for Spruce 1 (*p*<0.05) its explanatory power was weak (R^2^=0.05).

**Table 2.**
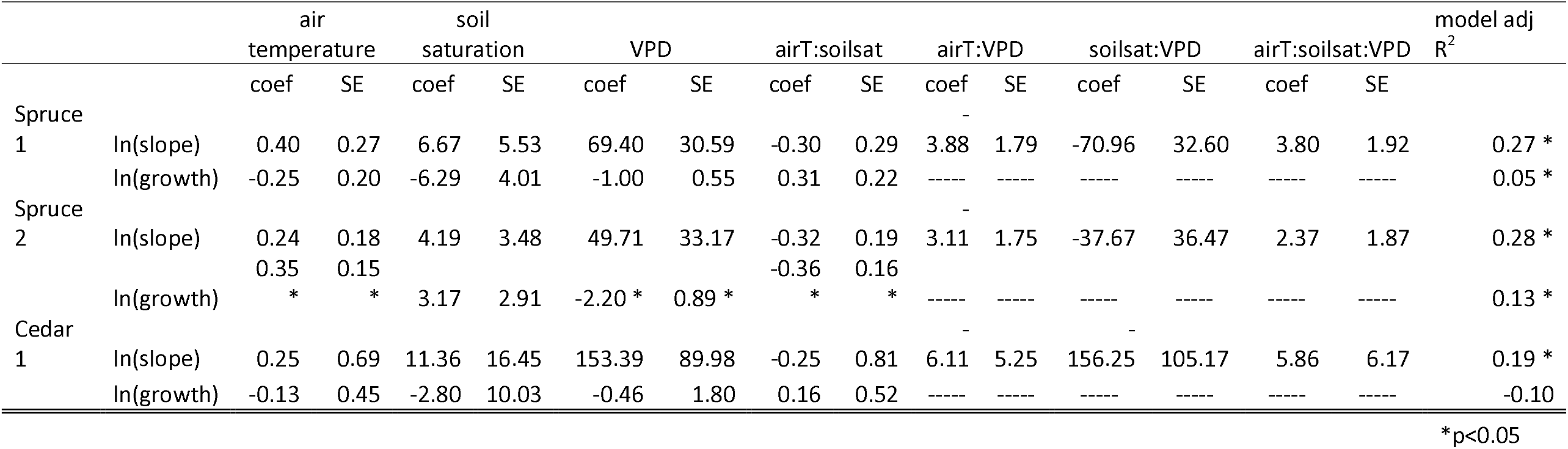
AIC selected models for linear regressions for the slope and growth of Spruce 1, Spruce 2, and Cedar against environmental variables. Each tree has a single model which includes all three environmental variables and their interactions. AirT = air temperature (°C), soilsat = soil saturation (mS cm^-1^), VPD = vapor pressure deficit (kPa), coef = coefficient, SE = standard error.

### CITs

We examined the statistical relationships between periods of growth and the ambient environmental conditions using conditional inference trees (Figure 5). This technique has the advantage that it not only partitions the growth response to local environmental conditions but also identifies specific values of these variables that can indicate a critical threshold of the response. Solar radiation, temperature, soil water content and VPD and precipitation were all found to be important, and distinct between the three trees (Figure 5). The rate of growth in Spruce 1 was significantly related to soil temperature with more growth happening when the soil is warmer than 17.5 °C. At cooler temperatures (below 17.5 °C), solar radiation became important with a critical value of 39.6 W m^-2^ splitting the response to two groups with higher growth rates with less light than with more light (Figure 5). However, if soil temperatures were above 17.5 °C, a second critical temperature threshold was detected at 22.5 °C and growth was again higher above this temperature. Below either of the two temperature thresholds (17.5 or 22.5 °C) solar radiation appears as a significant variable and in both cases lower light was associated with higher rates of growth. By comparison Spruce 2 and Cedar were most responsive to variables related to water, namely VPD and precipitation (Figures 5e & 5f respectively). Precipitation edged out VPD for Spruce 2, with higher growth rates above 0.19 mm of rain than below. Below this threshold VPD was identified as a significant driver of growth with a critical threshold of 0.37 kPa. The importance of the two parameters is reversed for Cedar, yet the break points and the direction of the response are quite similar: VPD was the strongest predictor and had a threshold of 0.31 kPa while precipitation was secondary, splitting at 0.12 mm. The critical VPD value for the rate of growth was slightly higher than that for the amount of growth (0.23 vs. 0.31).

**Figure 5.**
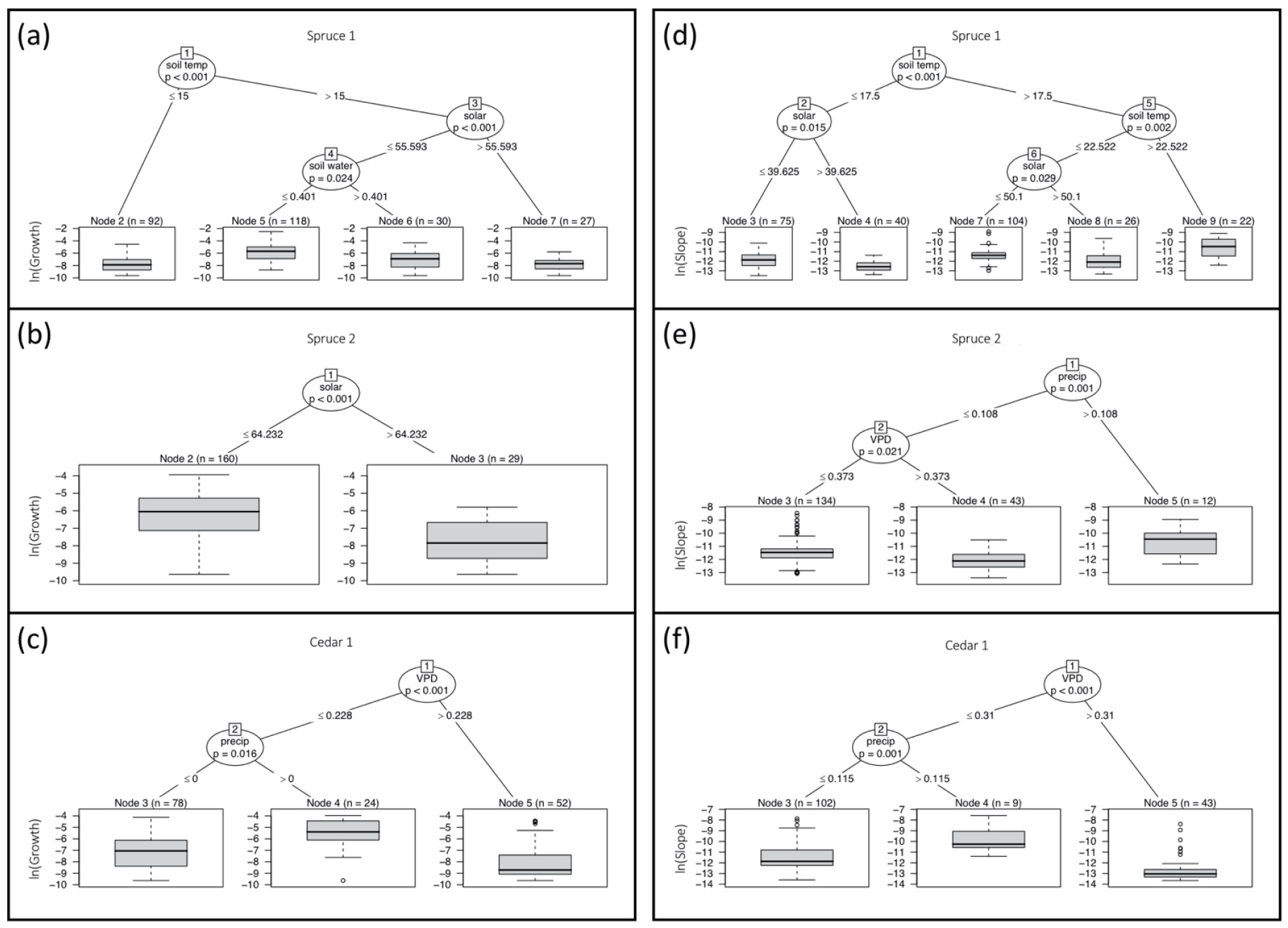
CITs partitioning the influence of environmental variables on total amount of log(growth) during HMM-identified growth periods (a-c) and on log(slope) across HMM-identified growth periods (d-f), calculated using values of the zero-growth model at the beginning and end points of each growth period. a) Spruce 1, growth. Soil temperature (soil temp) was the most important predictor, and solar and soil water were also identified as influential. b) Spruce 2, growth. Solar was identified as the most important predictor. c) Cedar, growth. VPD was the most important predictor, and precipitation (precip) was also identified as influential. d) Spruce 1, slope. Soil temp was the most important predictor, but solar was also identified as influential. e) Spruce 2, slope. Precip was identified as the most important predictor, with vapor pressure deficit (VPD) also identified as influential. f) Cedar, slope. VPD was the most important predictor, and precip was also influential.

More differences in relevant environmental variables and their break points were found in the amount of growth using the decision tree approach, especially for the spruces (Figure 5a, b). Soil temperature took the first position for Spruce 1 (threshold of 15 °C), followed by solar radiation (threshold of 55.6 W m^-2^), and then soil water (threshold of 0.4 m^3^ m^-3^, Figure 5a). Overall this is a simpler set of values than was found significant for the rate of growth of this tree, yet the most complicated of three total growth relationships observed. By contrast Spruce 2 had the simplest decision tree, experiencing more growth when solar radiation was below 64.2 W m^-2^ than when it was above (Figure 5b). Finally, the variables identified as important to cedar growth are the same variables identified as important to the rate of growth in this tree: VPD (0.23 kPa) and precipitation (>0 mm) and show more growth occurs when the VPD is low and it is raining (Figure 5c).

## Discussion

Dendrometers provide an accurate and precise record of short-(5 min) to long-term (annual) changes in stem diameter, giving a detailed signal of biological activity. This fine resolution allows comparison to environmental conditions that support growth from the hourly to yearly time scale. By streaming these data directly to the internet and adding co-located measurements of the environmental conditions, the system can provide highly useful, near real-time information and contribute to the “Internet of Nature” (Galle et al., 2019). Similar projects exist in Europe (e.g. TreeWatch (https://treewatch.net) and TreeNet (https://treenet.info)), and have proven to provide valuable information in built and natural environments. Expanded urban tree networks could allow homeowners or parks managers to optimize irrigation regimes to match watering to periods of rapid growth and/or water deficit (Ortuño et al., 2010), researchers to assess the impact of changing environmental conditions on carbon sequestration (Lada et al., 2019), and STEM educators to build lesson plans based on the scientific method and data collection for hypothesis testing (Sanders, 2012). Our system identified the duration of the growing season, and within these, periods of overall growth and dormancy. On finer time scales we show that growth (stem expansion) occurs primarily at night and depends on the interacting environmental variables relating to both energy gain and water use.

### Intra-annual diameter fluctuations

Understanding the links among the urban environment, tree biology and resultant ecosystem services requires the precise quantification of the relationships among these parameters. Generally, when temperatures are above freezing and physiological activity has a clear influence on stem diameter, contraction is observed during the day as transpiration moves water through the stem more quickly than it can be replenished from the soils via the roots (Goldstein et al., 1998; Steppe & Lemeur, 2004; Köcher et al., 2013). At night, stems generally expand as transpiration slows to a minimum and the hydrologic balance tips towards the absorption of water from the soils and stem refilling, providing the turgor pressure necessary for cell expansion (Steppe et al., 2015). At our site more than 59% of all stem expansion happens at night when temperatures are lower, humidity is higher, the vapor pressure deficit is lower, and soil moisture is abundant (due to regular irrigation at this site). These diel patterns in stem diameter provide a framework for the study of urban trees and forests and connect the observations to the underlying physiological processes that are themselves responding to environmental variation on similarly short-time scales.

To distinguish periods of growth from those of hydrologically driven fluctuations in stem radius, we used an existing model for partitioning the dendrometer data that “assumes zero growth during periods of stem shrinkage” (ZG model, Zweifel et al., 2016). This model limits its definition of growth to periods of time when new maximum stem radii are achieved and assumes stems cannot both shrink due to water deficit and grow due to cell expansion simultaneously, which has been shown (Lockhart, 1965; Steppe et al., 2006; Zweifel, 2016; Zweifel et al., 2016). We applied a HMM to the emitted states of the ZG model, which has the advantage of identifying blocks of time dominated by growth without requiring the dendrometer readings be monotonically increasing, so that even during a growth period growth need not be observed between every five-minute increment. The growth periods thus identified by the HMM help avoid pseudoreplication that would result from treating ZG “growth” time steps independently. The ZG and HMM methods are shown for two seven-day periods, one featuring consistent growth and the other showing a period of water limitation leading to stem shrinkage and the cessation of growth (Figure 4). During the six days in June, the ZG model identifies 64 individual growth periods, over 70% of which are 15 minutes or shorter; the HMM extracted three periods of growth over the same interval. With measurements being collected so frequently by the dendrometers, the HMM provides an effective way to identify separate growth periods for comparing growth rates to average environmental conditions, as discussed below.

We found both air temperature and soil and atmospheric moisture affected growth, contrary to what we expected from irrigated trees. In their native environment of boreal forests (Abrahamson, 2015), white spruce is thought to be commonly limited by both cool temperatures (Achuff & La Roi, 1977) and soil moisture (McGuire et al., 2010). Despite daily watering, the CIT for Spruce 1 in our experiment nonetheless identified low soil moisture along with colder soil temperatures as retarding stem growth. Previous studies have found that fluctuations in stem diameter are positively correlated with air (Ortuño et al., 2006; Devine & Harrington, 2011; Oberhuber et al., 2015; Dong et al., 2019), not soil temperatures. We suspect in our study, soil temperatures were buffered from short-term fluctuations in onshore winds and thus represent a more stable representation of the environmental temperature controlling growth of the stems inside these dense coniferous canopies. From the regression analysis, we conclude that spruce growth (the amount of expansion during the growth periods) is best explained by a model that contains information about air temperature, soil moisture, VPD and all of the interaction terms (Table 2). All of these variables are directly or indirectly related to environmental control of stem water status and have contributed to previous attempts to interpret dendrometer signals (Dong et al., 2019). While our best fit model can account for a significant amount of variation in the growth rate observations, it may have limited predictive capability, as more than 70% of the variance is unaccounted for. The reason for this lack of predictive power is unclear but may indicate a need to include information on substrate availability, time lags, or source/sink dynamics (Savidge, 1983, 1996, 2001; Dengler, 2001; Vaganov et al., 2011). Additional dendrometers spanning urban to rural ecosystems might enable further understanding of whether this decoupling from environmental conditions is a unique product of the urban environment.

We observed a similar pattern for cedar as described above for the spruce, yet the individual parameter estimates were quite different and indicate a stronger response to soil moisture and VPD. This difference may reflect the evolutionary history of this species. *Cryptomeria japonica* is an endemic monoecious conifer of Japan (Kimura et al., 2014) thought to have a distributional corresponding to moderate climatic conditions including annual precipitation of at least 1200 mm, annual temperature > 5.0 °C, (Tsukada, 1982; as cited by Kimura et al., 2014). The CITs suggest low VPDs are the strongest determinant of growth in this tree, with a critical threshold of 0.23 kPa (growth) and 0.31 kPa (growth rate). Previous work on stem shrinkage suggests the response is to an interaction between soil water (the source for stem water) and VPD (the driving force for water loss *via* transpiration) (Garnier & Berger, 1986; Hinckley & Bruckerhoff, 1975; Grossiord et al., 2017) and is consistent with our findings here for the constraints on stem growth. In general, trees from the Cupressaceae are believed to drought tolerant, due to the evolution of drought-resistant xylem and a general increase in the carbon investment in xylem tissue (Pittermann et al., 2012), yet drought sensitive since the tree rings of many members of this family faithfully record the occurrence of droughts (e.g. Sano et al., 2009; Buckley et al., 2017, 2018; B. Buckley pers. com.). Previous work on cedar (Abe & Nakai, 1999; Abe et al., 2003), shows a detailed response of xylem development to stem water status suggesting a cedar is highly sensitive to water availability. Just as with spruce, the linear model for cedar had a greater capacity to account for variation in the rate of growth than in the magnitude of growth, yet even the rate model only accounted for 19% of the variance. Clearly, our ability to predict short-term fluctuations in the rates and amounts of growth requires further study, particularly in the urban environment, and a more complete understanding of the physiological controls of cambial activity. We also note the potential influence of lag times and time accumulation effects of climate on tree growth, which could decouple the high-resolution measurements of growth from the environmental drivers on these short time scales. Thus on the whole, we find only partial support for our hypothesis that growth in urban irrigated trees would be more strongly controlled by environmental factors related to energy gain (temperature and solar radiation) than by factors related to water use (precipitation, VPD and soil moisture).

### Seasonal growth trends

Tracking the phenology and climatic response of trees in urban settings is critical to understanding both the ecosystem services urban trees provide and how global climate change might affect these services. Vegetation phenology, and particularly tree growth, is often studied using remote sensing surveys of spectral reflectance (e.g. Ren et al., 2018), yet these studies can be challenging in urban settings due to the highly heterogeneous urban landscape and the relatively large spatial scales and infrequent flyover times of satellites (Melaas et al., 2016). Dendrometers can quantify both the short and long-term phenology of tree growth allowing urban arboriculturists to gain insights into the specific environmental conditions stimulating and retarding tree growth, affording the opportunity to initiate tree care when it will be most effective.

The onset of growth is likely triggered by warming temperatures (Begum et al., 2013), and changes in daylength (Oribe & Kubo, 1997; Chang et al., 2020). In the urban setting studied, we find evergreen tree growth begins in late April and continues through the spring, summer, and fall, ending in late September (spruce) or late October (cedar). Spruce growth is coincident with a daylength of 13hr, 20 min and an accumulated growing degree day of 69.7 °C, whereas the initiation of radial expansion in the cedar is associated with longer days and much higher thermal accumulation (13hr, 45min and 160.6 °C), possibly due to bark thickness which may affect the relative rate of warming and the breaking of cambial dormancy (Oribe & Kubo, 1997). As urban environments are typically warmer than their rural counterparts, they may have an earlier start to the growing season (Yang et al., 2020). A monitoring system along an urban to rural gradient could provide additional insights into how urban environmental conditions affect tree growth.

Detecting the onset of radial growth may go unnoticed as it begins well in advance of leaf growth of the more common deciduous trees native to our study region and even before leaf and stem elongation in many evergreen species (Oribe et al., 1993; as cited in Oribe & Kubo, 1997). In temperature ecosystems, radial growth continues throughout the spring, summer, and early fall and demonstrates a clear seasonal pattern, related to cold hardiness and surviving the winter (Catesson, 1994). Cold temperatures, a decreasing temperature trend, photoperiod, and changes in light quality (Chang et al., 2020) all cue the end of growth and cold hardening. After a brief resting phase when cell division is not possible, the cambium will remain in a quiescent stage (Catesson, 1994), capable of cell division once environmentally favorable conditions return. Dendrometers provide the opportunity to observe and even quantify these various stages of activity; placing dendrometers in urban settings enables study of how the built environment might alter the growth and cold-response of trees. Without real-time dendrometer data, the evergreen growth form of our study trees would have made the phenological timeline very difficult to detect and thus we suggest that networks of dendrometers in urban areas would also provide an effective tool for arboriculturist to monitor growth and precisely time pruning, irrigation, and nutrient supply.

While urban areas are responsible for more than 70% of CO_2_ emissions (Global Energy Assessment Writing Team, 2012), the contribution of urban trees to directly offset this is not insubstantial. Previous work by Nowak and Crane (2002), estimated trees in New York City alone may store as much as 1.2 million tonnes of carbon, with an annual rate of net carbon sequestration of more than 20,000 tC yr^-1^. Using a valuation of $20.3 tC^-1^ (Fankhauser, 1994), Nowak and Crane (2002) estimated that the 700 million tonnes of carbon stored by urban trees in the US had a value of more than $14 billion in 2001. The four small trees in our study gained a total of 45.4 kg of above ground dry mass, thereby sequestering 22.7 kg C, or 83.3 kg of CO_2_ during the duration of the study (July 2018 August 2019, the time when the Spruce 1 sensor saturated) (Table 1). Adding 30% to conservatively account for changes in root mass, increases the amount of carbon sequestration to 29.5 kg of C or 108.3 kg of CO_2_. Estimates like these, based on precise data from dendrometers in a variety of urban settings, would help refine our estimates of carbon storage provided by urban trees. As robust estimates of carbon storage and cycling become more important in light of climate change, land use change, and urbanization, networks of dendrometers in urban settings can play an important role providing urban foresters and arboriculturist a reliable tool for monitoring, quantifying, and valuing the urban contribution to biological carbon sequestration.

### Conclusions and further opportunities

We installed high-resolution dendrometers in an urban setting and streamed the data to a project web page in near real time. The resulting time series provides insights into the growth of specimen trees growing in a residential area. From this time series were able to assess the relationship between fluctuations in stem diameter and the ambient environmental conditions, the phenology of stem growth, and the magnitude of growth and carbon sequestration. Although we had hypothesized that stem growth would respond most strongly to light and temperature, our results demonstrate that these trees have a discernible response to environmental water relations (soil moisture, precipitation and VPD) despite being irrigated. We also introduce new analytical techniques, HMM and CITs, to interpret established ones (multiple linear regression models, and the zero growth model). Overall, we suggest CITs as a useful way to search for important environmental variables and thresholds which can then be used to test more specific hypotheses related to the environmental control of stem growth and hydrology. Furthermore, as compared to the linear models, the decision trees are more sensitive to unique subsets of the time series and thus can identify important relationships that the overall fit of the linear model might miss. We show that three modest ornamental trees sequestered 22.7 kg C (or 83.3 kg of CO_2_) between July 2018 and August 2019. We further demonstrate that the stem expansion is primarily a nocturnal phenomenon while stem shrinkage corresponds to the daylight hours. Although there were some important differences among our three trees, there are obvious general instantaneous relationships with temperature, VPD and precipitation, suggesting water relations control growth. Detailed information like this is rare in urban environments and yet facilitates a deeper understanding of dynamics that is useful to professionals involved in urban ecosystem management.

Dendrometers can be a powerful tool in urban settings and can provide care and maintenance insights for land owners and land managers in urban areas where trees provide important ecosystem services. With the data streaming to the web, would allow easy access to data by classrooms and nature centers, providing a more proximate connection for urban residents to the natural world (Kitchin, 2014; Galle et al., 2019). Fluctuations in stem diameter happening over remarkably short time scales and tied to the local environmental conditions surprise people and bring tree biology to life. Sharing these data with the public at nature centers and during other outreach opportunities, or working with teachers in a K-12 educational setting, has always piqued curiosity and stimulated deeper thinking and inquiry in STEM topics. These real time data sets also provide an opportunity for teaching the scientific method, beginning with observation then generating hypotheses, data analysis, hypothesis testing and finally interpretation and the ability to begin the process again immediately, and with new information. Live-streaming tree growth data facilitates an even deeper understanding and appreciation of the biological activity of trees and the services they provide in urban environments and thus can be a powerful tool connecting urban social and ecological systems.

